# Identification of a mimotope of a complex gp41 Human Immunodeficiency VIrus epitope related to a non-structural protein of *Hepacivirus* previously implicated in Kawasaki disease

**DOI:** 10.1101/2024.06.26.600771

**Authors:** Hakimuddin Sojar, Sarah Baron, Mark D Hicar

## Abstract

**Background:** We have previously isolated a highly mutated VH1-02 antibody termed group C 76-Q13-6F5 (6F5) that targets a conformational epitope on gp41. 6F5 has the capacity to mediate Ab dependent cell cytotoxicity (ADCC). When the VH1-02 group C 76 antibodies variable chain sequence was reverted to germline (76Canc), this still retained ADCC activity. Due to this ability for the 76Canc germline antibody to functionally target this epitope, we sought to identify a protein target for vaccine development.

**Methods:** Initially, we interrogated peptide targeting by screening a microarray containing 29,127 linear peptides. Western blot and ELISAs were used to confirm binding and explore human serum targeting. Autoimmune targeting was further interrogated on a yeast-displayed human protein microarray.

**Results:** 76Canc specifically recognized a number of acidic peptides. Meme analysis identified a peptide sequence similar to a non-structural protein of *Hepacivirus* previously implicated in Kawasaki disease (KD). Binding was confirmed to top peptides, including the *Hepacivirus*-related and KD-related peptide. On serum competitions studies using samples from children with KD compared to controls, targeting of this epitope showed no specific correlation to having KD. Human protein autoantigen screening was also reassuring.

**Conclusions:** This study identifies a peptide that can mimic the gp41 epitope targeted by 76C group antibodies (*i.e.* a mimotope). We show little risk of autoimmune targeting including any inflammation similar to KD, implying non-specific targeting of this peptide during KD. Development of such peptides as the basis for vaccination should proceed cautiously.

## Background

The creation of a successful human immunodeficiency virus (HIV) vaccine continues to be a public health priority^1^. A large effort has been focused on discovery and characterization of broadly neutralizing antibodies (bnAbs) ^2, 3^. Many bnAbs are highly mutated, but increased levels of mutations can be stochastic and do not predict neutralization. The monoclonal antibody (Ab) 76-Q13-6F5 (6F5) is highly mutated (83% homologous to predicted heavy chain germline), and has the capacity to mediate Ab dependent cell cytotoxicity (ADCC) ^4^. The 6F5 epitope encompasses areas in both heptad repeats of gp41, mapping by alanine scanning mutagenesis to amino acids (AA) R557, E654 and E657 of reference sequence HXB2, just proximal to the membrane-proximal external region (MPER-underlined) ^5^. Three other Abs (76-Q11-4E4, 76-Q7-6F11 and 76-Q7-7C6) used VH1-02, competed for the 6F5 binding, and were also shown to target E657 AA (bold) (AA 652-667: QQEKN**<u>E</u>**QELL<u>ELDKWA</u>) ^5, 6^. We grouped these into an epitope targeting group termed 76C Abs.

Serum from HIV long-term nonprogressors contained significantly higher levels of 76C Abs in comparison to HIV infected persons with comparable viral loads. Due to this correlation with non-progression, further studies were done by creating a 76C group ancestor (76Canc) Ab utilizing the unmutated germline heavy variable chain from VH1-02. From exploring the derivation and possible cross-reactivity of 76Canc, we discovered this ancestor Ab also has significant functional ADCC activity ^4^.

Development of Abs utilizing VH1-02 gene segments after vaccination is well studied, as this is used in VRC01, one of the most bnAbs ^7, 8^. VRC01 is highly mutated, with V-gene region AA predicted mutations of 42% in the heavy chain and 28% in the light chain. The recognition of the CD4 binding site is predominantly driven by this heavy chain ^7^ and related Abs rely on similar structures ^8, 9^. Neutralization was maintained when VRC01 framework mutations were mutated to ‘near’ germline ^10^, but unmutated common ancestors of these Abs don’t interact with native trimers, creating further challenge for vaccination strategies based on the concept of stimulating the naïve Ab repertoire to generate a HIV bNab response ^8, 11–14^.

As we have shown that Abs related to 6F5 correlate with non-progression and that germline use of VH1-02 in 76Canc can support anti-HIV functional ADCC, we propose a vaccine strategy to create such 76Canc-like Abs. Unfortunately, a number of studies utilizing gp41 constructs, including trimeric forms, have been relatively unsuccessful ^15^. In this study, we sought to discover a protein target that could be recongnized by 76C group Abs to be used in future immunization studies.

## Methods

### Enrollment

Plasma samples from febrile children including KD subject samples (UBKD) and associated clinical information were collected under approval of the UB IRB STUDIES-00000126, 00002824 and 00005262 with funding support by the Wildermuth Memorial Foundation as previously described ^16^. Additional serum samples (30 complete KD subjects with pre-intravenous immunoglobulin (IVIG) treatment, post-IVIG, and convalescent samples) were obtained through the Pediatric Heart Network and stored in the Kawasaki Disease Biorepository (KDB) at Boston Children’s Hospital (IRB X10-01-0308) which were collected for a prior study ^17^. Statistical analysis was performed using GraphPad Prism 9 and groups were compared with Wilcoxon ranked sum tests.

### Serum antigen targeting screening

Serum samples were provided to CDI laboratories to screen on the HuProt array. The HuProt array is a yeast-derived expression library of 23,059 human proteins. These targets are duplicated on the screens and binding is normalized to background binding and calculated per company’s protocols. Specific Abs were screened per company protocols on the PEPperCHIP^®^ Human Epitome Microarray, containing 29,127 linear peptides printed in duplicate. The peptide content was based on all linear B-cell epitopes of the Immune Epitope Database with the host “human” and was further complemented by all epitopes of the most common vaccines.

### Meme analysis

The top 65 Ab targets identified on the PEPperCHIP^®^ Human Epitome Microarray (>200 fluorescence units threshold) were uploaded to the MEME tool (http://meme-suite.org//tools/meme). The MEME pre-settings were a maximum of one motif per each sequence with maximum total 5 different motifs, as well as a minimum motif length of 4 AA and threshold of E < 5.0e-002.

### Peptide ELISA and Characterization

Peptide ELISAs proceded as previously described ^18^ with the following adjustments: peptides were dissolved in 50% DMSO in PBS and coated at 10 ng/well of peptide and incubated overnight at 4°C on a rocking platform prior to assay. For biotinylated Ab competition ELISAs, Ab biotinylation and ELISA was performed as previously described ^4^. Peptide characteristics (isoelectric point, charge at pH 7 and hydrophilicity) were calculated with online calculator (Bachem.com) with N-terminal -H and C-terminal -OH.

### Protein Binding ELISA: Confirmation of autoantigen targeting

For Western blotting and ELISA assays, human glutaredoxin 3 (GLRX3, catalog # TP302731) and human Tropomodulin1 (TMOD1, catalog # TP301134) were obtained from OriGene Technologies Inc, Rockville, MD. TMOD1 human recombinant isoform 1 (NP_003266.1) and GLRX3 isoform 1 (NP_006532.2) were used in BLAST analysis. Recombinant protein ELISAs proceded as previously described ^19^ with the following adjustments: proteins were plated at 10 ng/well overnight at 4°C, for GLRX3, 1% BSA was used as diluent, and for TMOD1, 7.5% FBS in PBS was used as diluent.

### Slot blot analysis

Peptides were transferred onto blotting membrane using Bio Dot Microfiltration system (Bio Rad Chemical Cat#170398) according to the manufacturer’s instructions, blocked with 1% BSA in pH 7.5 Tris-Buffer saline for 1 hr at room temperature. After rinsing, primary Ab was diluted in 1% BSA in Tris-Buffer saline pH 7.5 and incubated overnight at 4°C. Blot was washed (3 × 10 minutes) with gentle agitation. Secondary Ab (Alkaline phosphatase -conjugated anti-human IgG, Southern Biotech, Birmingham, Al) was added in 1% BSA in Tris-Buffer saline pH 7.5 and incubated for 1 hour at room temperature with gentle agitation. Blot was then washed three times in Tris buffer saline pH 7. Bands were visualized with Alkaline phosphate substrate NBT/BCIP (Thermo Scientific, Grand Island, NY).

## Results

As the epitope targeted by the group 76C Abs is conformational, further definition of this region can be facilitated by isolating peptides that can replicate such discontinous conformational epitopes; or so called mimotopes ^20^. We interrogated the PEPperCHIP® Human Epitome Microarray, covering 29,127 linear peptides, to search for possible mimotopes. The peptide content was based on all linear B-cell epitopes of the Immune Epitope Database with the host “human”, and was further complemented by all epitopes of the most common vaccines.

On library screening using 76Canc, which has the unmutated VH1-02 segment, the most significant binding was against a number of negatively charged peptides from glycinin (*Arachis hypogaea*) with the consensus motif EYDEDEYEY (**Figure 1**). Most of the top hits were highly acidic with the average isoelectric point of the top 20 being 3.05 with charge at neutral pH of -6.84. This is not surprising since the 76 group C epitope is in an acidic hydrophilic region in the carboxy-terminal heptad repeat (Hxb2 gp160 reference AA 652-667: QQ**<u>E</u>**KN**<u>E</u>**QELLELDKWA; bolded/underlined resolved by alanine scanning mutagenesis^5^). Numerous possible human pathogen motifs were identified with many of these being negatively charged. A number of human peptides enriched for negatively charged acidic AA were also readily recognized from the cerebellar degeneration-related antigen 1, Major Centromere Autoantigen B, and coagulation factor VIII precursor.

**Figure 1:**
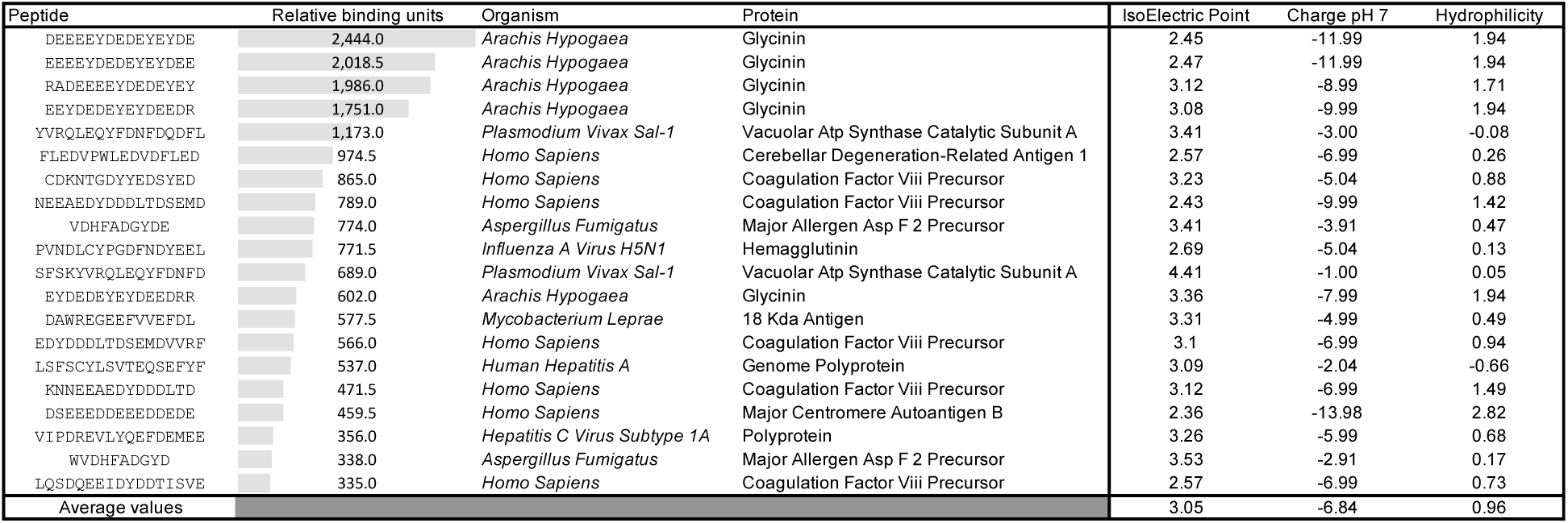
Top 20 peptides recognized by 76Canc. Relative binding units are shaded in comparison to zero, which is the normalized background. Peptide characteristics (isoelectric point, charge at pH 7 and hydrophilicity) were calculated using the online calculator (Bachem.com).

No HIV-related peptides showed significant binding activity (detailed in **supplemental table 1**), consistent with lack of gp41-derived peptide binding in prior studies ^5^. This included six HIV peptides that overlapped with the 76C group E657 motif (red text, **supplemental Table 1**) all which had minimal binding (<50 relative binding units) including the very acidic peptide EELKQLLEQWNLVIGFL (ie 3.95). As HIV and coronaviruses (CoVs) are both type 1 fusion proteins, it is plausible there is some cross-reactivity between Abs that may target a structural domain on the fusion proteins (HIV envelope and CoV Spike). Peptides derived from SARS CoV were included in the peptide screen and showed no appreciable binding, including the Spike S2 peptide PLKPTKRSFIEDLLF, which is homologous to the 76C group epitope on gp41.

### Meme analysis

To explore consensus targets, a MEME analysis of all peptides with a spot intensity of >200 fluorescence units (top 65 hits, details in supplemental table 1) was performed (**Figure 2**). The top motif exhibited a very high statistical significance of E = 5.3e-083 with contributions from 10 of 65 top hits and a motif length of 17 AA. This motif mainly originated from various similar *hepatitis C virus* (HCV) peptides. Due to the uncommon epitope length, it’s possible these peptides could replicate a conformational epitope. It’s also possible the main motif was based on a shorter acidic portion of the C-terminal, as the second motif (Figure 1, REVLYxxFDEM) was a shorter sequence within the first motif. It appears unlikely the FDEM sequence alone is targeted as there were 64 HCV FDEM containing peptides in the screen, but only 15 with spot intensity of >200 fluorescence units (see **supplemental table 1**).

**Figure 2:**
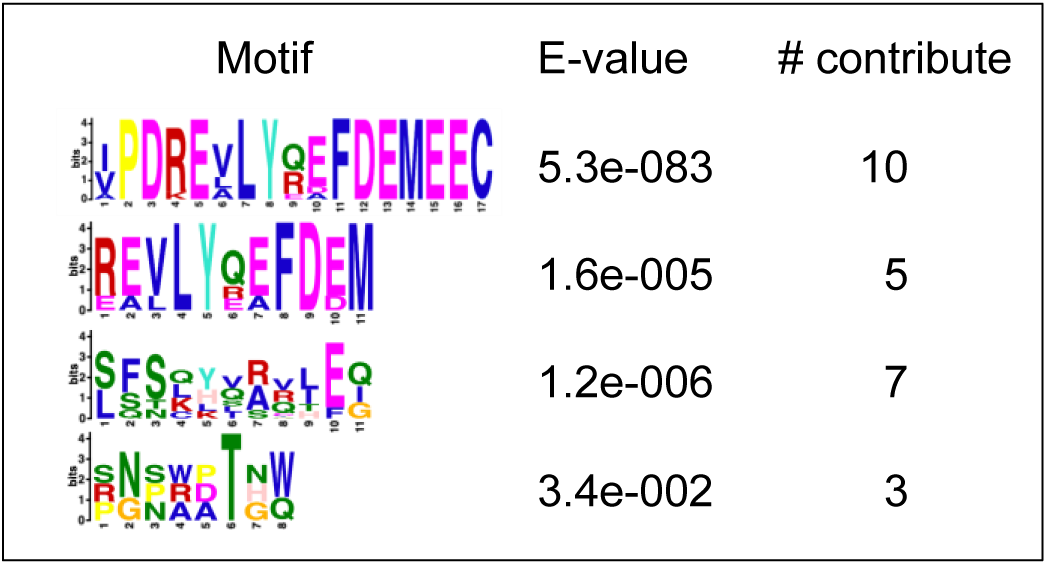
Motif Meme analysis of top 65 peptides recognized by 76Canc. The top 65 identified peptides were analyzed using meme analysis, http://meme-suite.org//.

### Confirmatory binding

Five peptides that reflected top peptide hits (**Table 1**) and the meme analysis were produced and compared to peptides from a number of pathogens of interest, and an acidic peptide from *Plasmodium falciparum* that did not show appreciable binding on the peptide microarray. On ELISA assay, biotinylated Abs of 76Canc, 6F5 and 6F11, bound all five top-hit peptides over twice the background of the control Ab (**Figure 3A**). Notable other targets showed specific binding from the top 65 hits (outer membrane protein of *Neisseria meningitidis* and AA permease of *Staphylococcus aureus*). A collection of acidic peptides (Table 1: 8, 9, and 10) and the blank well (50% DMSO only) were negative (peptide data 9 and 10 not shown). A slot blot assay was performed and confirmed binding to a number of these peptides, roughly corresponding to the level over background in ELISA results (**Figure 3B**, **Table 1**).

**Figure 3:**
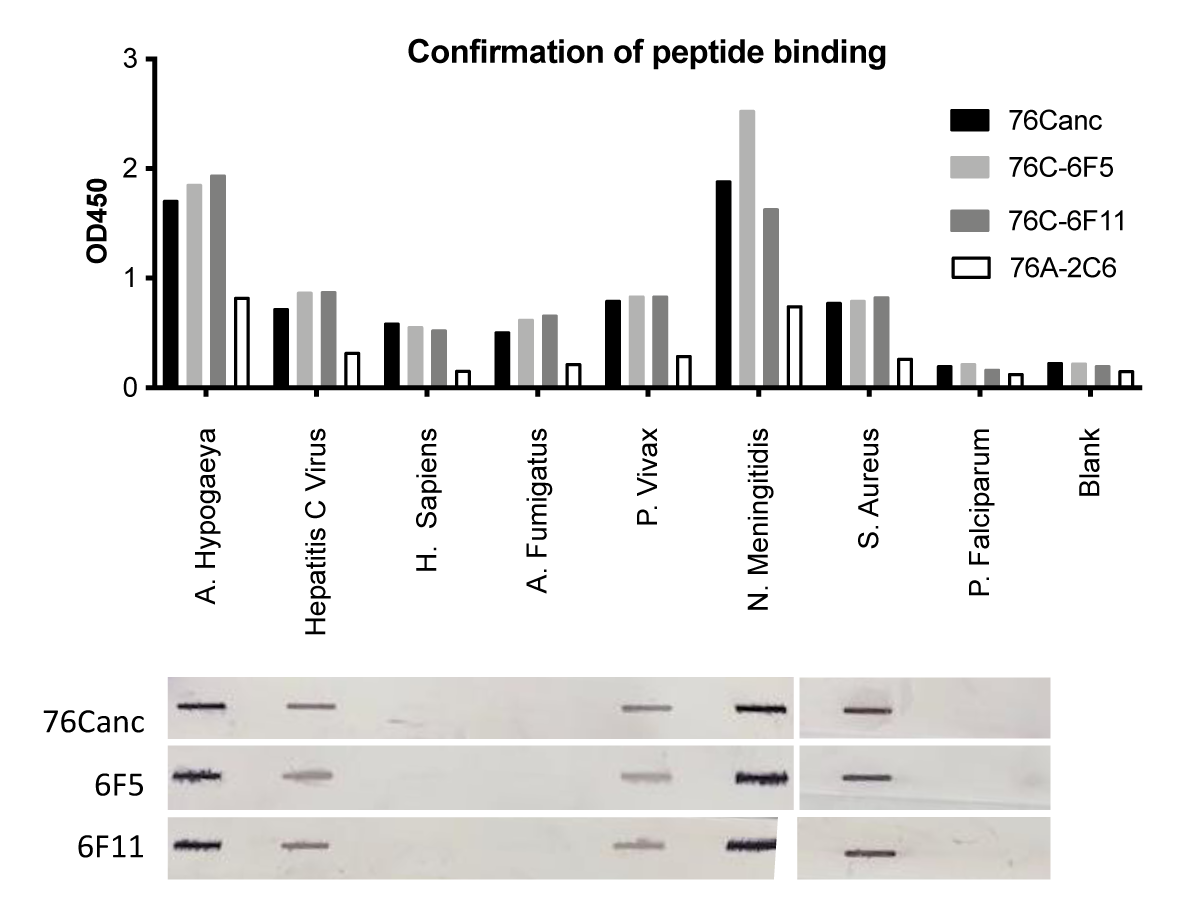
Confirming binding to peptides representing meme analysis. Peptides representing top hits and various controls from peptide screen with confirmation by A) ELISA assay using comparable parameters to original peptide screen (5 ug/mL of Ab) and B) slot blot Western blot (results shown all from a single blot, image was arranged to align to the ELISA data).

**Table 1:**
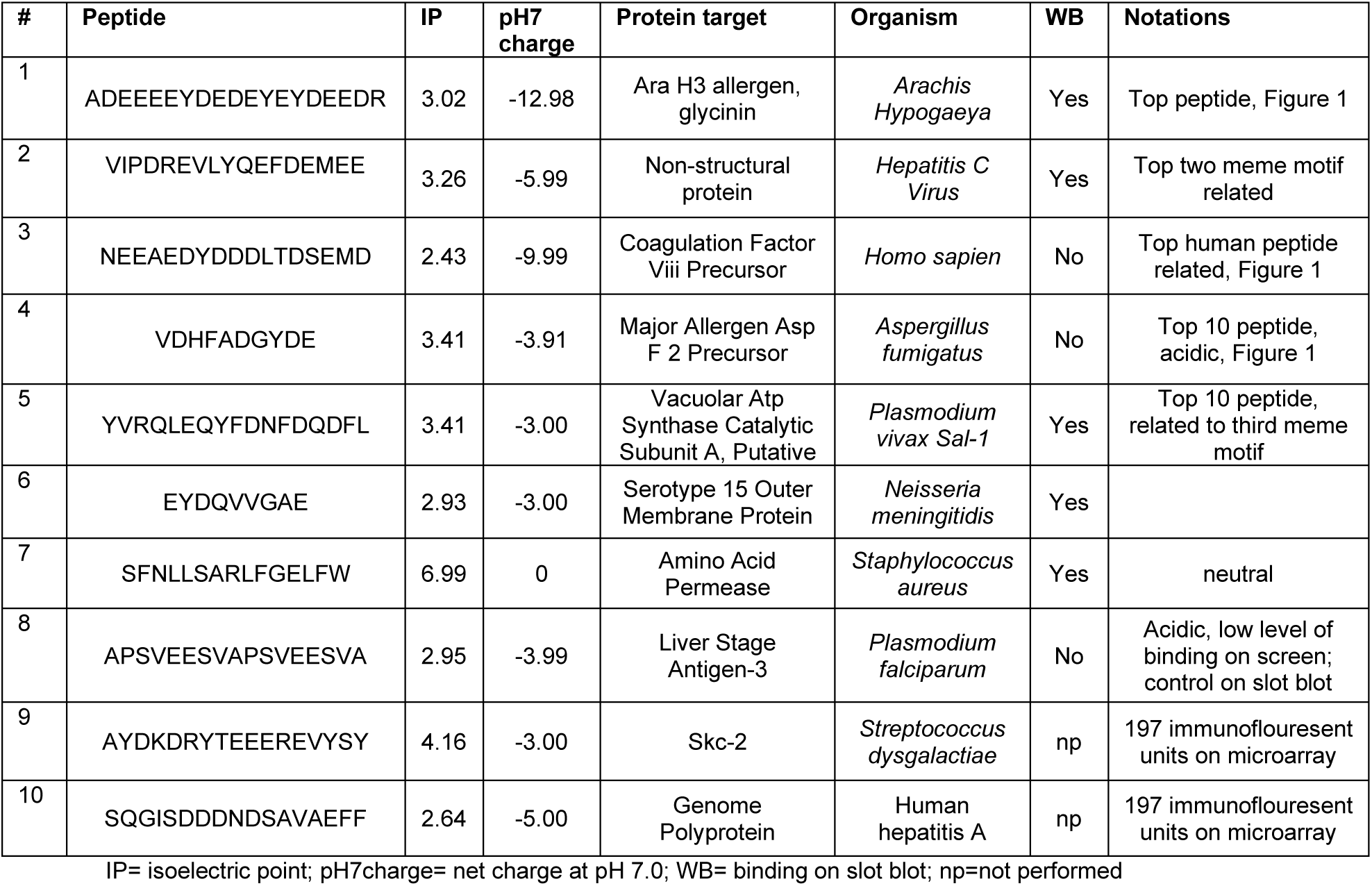
Selected peptides from peptide microarray screen produced for confirmation.

The microarray contains over 5,000 peptides from human proteins. A number of human peptides were in the top 65: Cerebellar Degeneration-Related Antigen 1, Coagulation Factor Viii Precursor, Major Centromere Autoantigen B, Kinesin-Like Protein Kif11, 78 kDa Glucose-Regulated Protein, Glutamate Decarboxylase 2, Calcium Channel, Alpha 1A Subunit Isoform 3, Heat Shock Protein 90Kd, DNA-Directed RNA Polymerase Iii Subunit, Rpc1, Trinucleotide Repeat Containing 6A and Isoform CraB Envoplakin. We did express the top human peptide (Table 1, peptide #3), which showed binding over twice background on ELISA (Figure 3), but was not shown to bind on slot blot analysis.

### Hepatitis C virus (HCV)-related peptide

The HCV-related peptide identified herein is similar to a recently identified peptide advanced to diagnose Kawasaki disease (KD) ^21^ KD4-2H4 KPAVIPDREALYQDIDEMEEC. This peptide was derived from a non-structural protein of HCV. KD is a vasculitis of children thought to be related to an infectious disease ^22, 23^. Despite an extensive history of studies attempting to associate an infection with KD, the cause of KD remains unknown^24^. In prior published studies using KD4-2H4, the specificity of binding was assay dependent, as there appeared to be binding by immunohistochemistry, but high concentrations of Abs were needed to show appreciable binding in ELISA (>1ug/mL)^21^.

We compared binding of KD4-2H4 to Peptides #1-3 from **Table 1**. We show that the binding of 6F5 and 6F11 readily recognizes all of these peptides with diminished binding by the 76Canc ancestor compared to the HIV 6F5 and 6F11 Abs (**Figure 4)**. Notably, reviewing the history of subject 10076 from whom these Abs were originally derived ^5, 6^, it should be noted this subject did not report a peanut allergy and were repeatedly negative on HCV testing.

**Figure 4:**
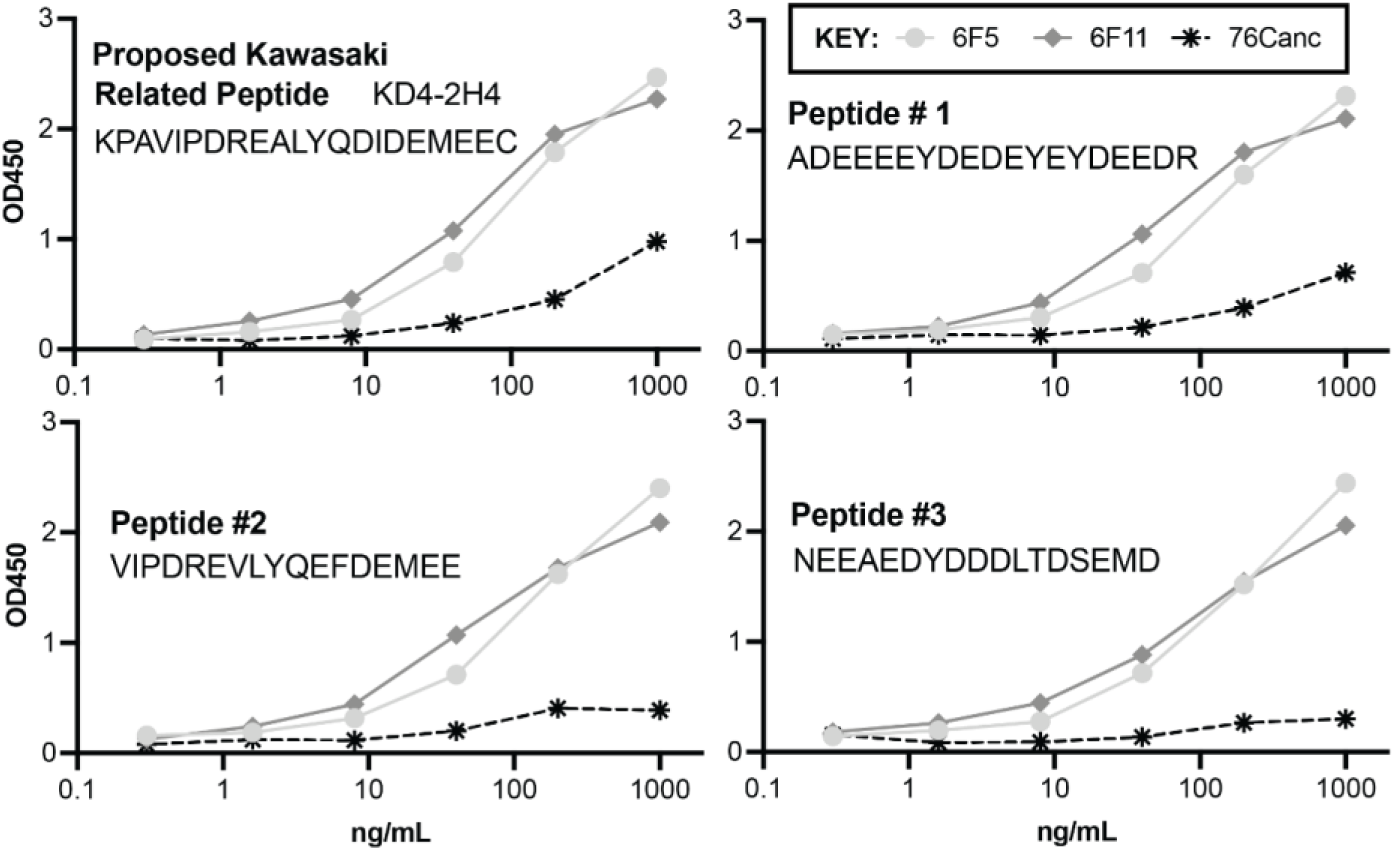
76C antibodies all recognize Hepatitis C Virus-related peptide implicated in KD. ELISA binding to KD4-2H4 and the top three peptides on our screen were performed using 76Canc (starred), 6F5 (light grey circle) and 6F11 (grey diamond).

### Clinical correlations

Although the cause remains unclear how aneurysms form during KD, autoantibodies (autoAbs) targeting is one of the proposed mechanisms ^23, 25, 26^. Since these 76C group Abs readily recognize KD4-2H4, we sought to assess if there was a correlation in 76C group Abs to KD. The 6F11 Ab was biotinylated and serum from a cohort of children with KD and febrile controls were used. Competitions of serum to 6F11 binding to KD4-2H4 showed no differences (**Figure 5**, mann-whitney p = 0.44) between KD and febrile controls. We additionally assessed a cohort of 30 children with complete KD, with serial pre-IVIG, post-IVIG, and convalescent samples, as previously described^27^. Overall, there was not a significant increase in KD4-2H4-targeting Abs that occurred in convalescent KD samples (pre-IVIG vs convalescent sample mann-whitney p value 0.77). After IVIG administration, there was no appreciable dilutionary effect in the majority of individuals, implying most IVIG formulations already contain Abs that would similarly bind this antigen. In subgroup analysis comparing those with elevated coronary artery Z scores, there was no overall difference between those with or without aneurysms for the pre-IVIG, post-IVIG and convalescent comparisons (mann whitney p = 0.71, p= 0.41, p = 0.24).

**Figure 5.**
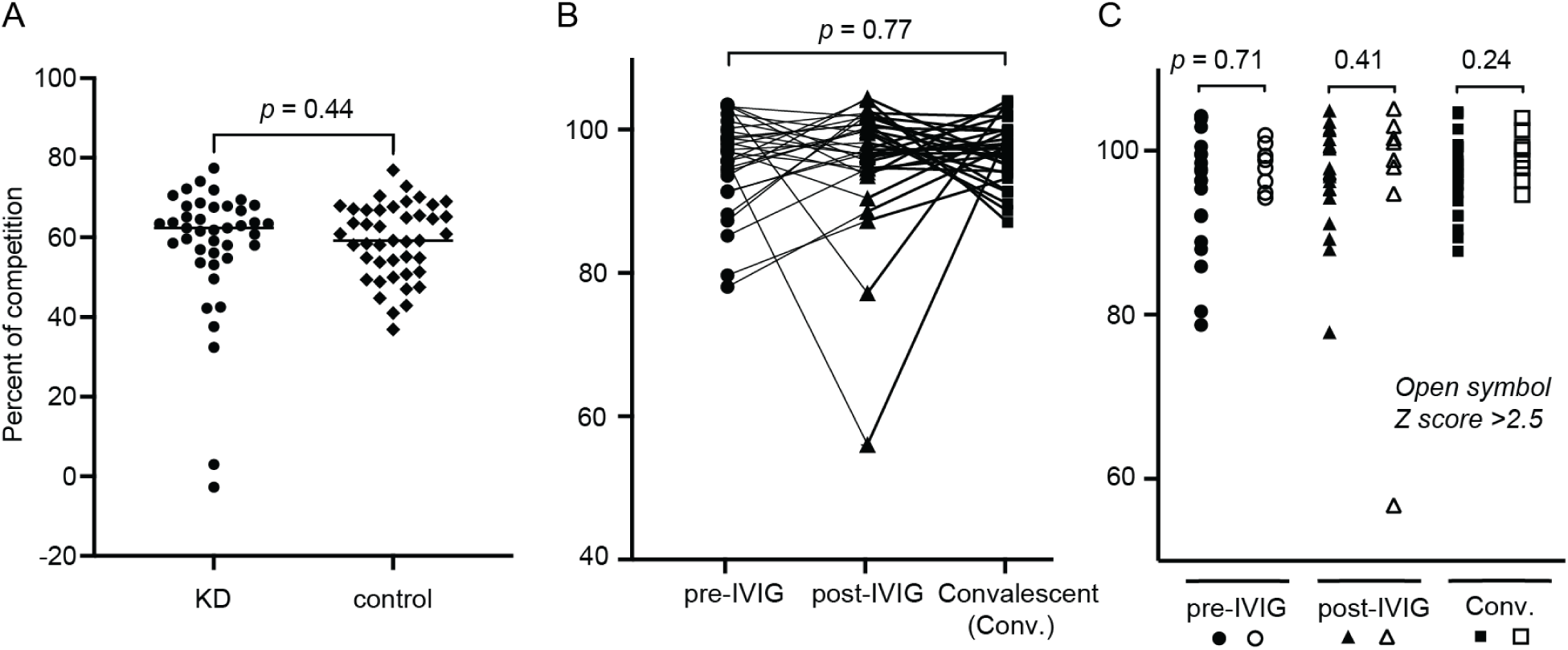
Humoral immune targeting to Hepatitis C Virus-derived peptide does not specifically identify children with KD. A) Serum at 1:200 was used to compete against biotinylated 6F11 binding to KPAVIPDREALYQDIDEMEEC. This was normalized to background negative competition wells as reading was 0% competition in KD (circle) and controls (diamond); B) Immune targeting was assessed in serial samples (pre-IVIG -circle, post-IVIG -triangle, convalescent -square) from 30 individuals with KD. C) Boston scoring for coronary artery aneurysms was used to define Z scores > 2.5 (open symbols) as previously published ^27^.

### Autoimmune assessment

We utilized the HuProtTM library (CDI Labs), a yeast derived expression library consisting of 23,059 purified human proteins to further assess potential autoimmunity. We compared binding of 6F5 Ab with the 76Canc (**Figure 6**). 6F5 (gray dots) showed a number of cross-reactions of unclear significance, the highest of which was Glutaredoxin 3 (GLRX3). Reactions shown on the peptide array interrogation were not replicated. Overall, 76Canc had generally less autoantigen binding than it’s more mature relative. The binding to Tropomodulin 1 (TMOD1) was a notable exception on this screen. GLRX3 is a fairly acidic protein, with a theoretical PI of 5.31 and containing 14.6% acidic AA. TMOD1 also had a theoretical PI of 5.01 containing 16.7% acidic AA.

**Figure 6:**
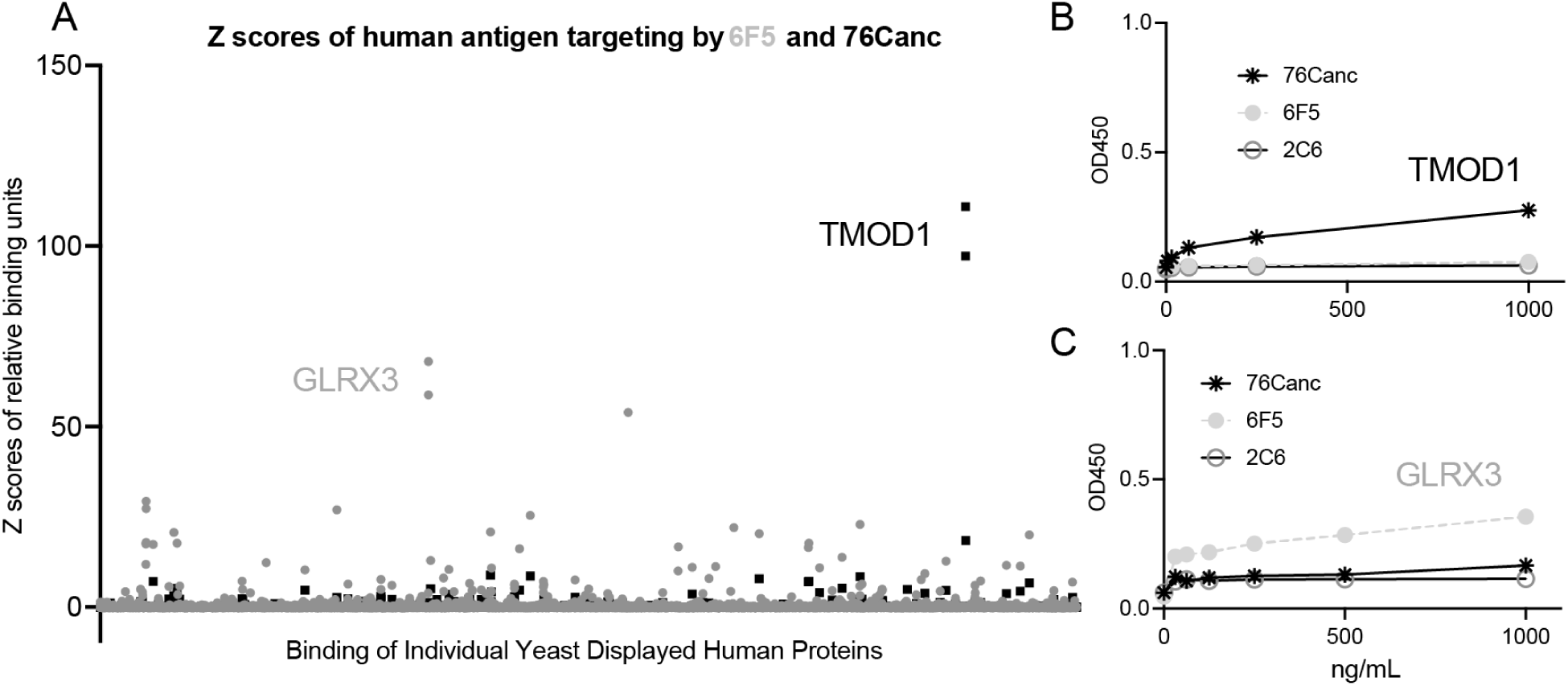
Binding to yeast displayed human proteins. A) Results of the full library interrogation for 23059 *S. cerevisae* expressed and purified human proteins (HuProt library, CDI labs) are displayed, for 76Canc (black) and 6F5 (gray). ELISA confirms low level binding of B) 76Canc to TMOD1 and C) 6F5 to GLRX3.

ELISA testing on recombinant GLRX3 and TMOD1 showed similar modest binding patterns as shown in the array. Western Blot analysis showed inconsistent resolution of binding (not shown). BLAST alignment did not reveal significant homology between TMOD1 and GLRX3, but did show portions of TMOD1 (AA 26-38) ELRTLENELDELD and GLRX3 (AA 231-243) KAPKLEERLKVLT that independently aligned with portions of the 76C group epitope. This further suggests the acidic nature of these epitopes may be contributing to this cross-reactivity.

## Discussion

In this study, we initially sought to discover a protein target that could potentially replicate the epitope (*i.e.* a mimotope) targeted by the 76C group Abs to be developed for future immunization studies. Surprisingly, a peptide identified by our anti-HIV Abs was highly similar to a peptide implicated in KD. We had initial concern in developing this peptide into a vaccine candidate due to the published findings with KD.

### Relationship to Kawasaki disease (KD)

It is unclear how Abs targeting this peptide relate to KD. There are no direct sequnceing studies that show any *Hepacivirus* member is related to KD. New PHIP-seq ^28^ approaches have also failed to show an association^29, 30^. Notably, prior studies have attempted to link CoVs as the cause of KD ^31^, but as reviewed, there were not significant targeting of CoV related peptides in our screen. Also, in our own prior studies comparing KD to febrile controls, we did not note any specific differences in targeting the Spike proteins of various CoVs, including SARS-CoV-2^27^.

The Abs that originally identified the KD4-2H4 peptide were derived from plasmablasts. We have shown that KD children have similar plasmablast to children responding to an infection ^16^, so conceptually this is a plausible approach. Antigen specificity has been shown when peripheral plasmablasts levels peak, usually 5-10 days after antigen challenge ^21^. In our prior study, the peak of plasmablasts in KD was on day 5 of fever. It’s reported that the Abs that originally identified the KD4-2H4 peptide were derived from plasmablast roughly two to three weeks into fevers. If this KD4-2H4 peptide was identified by such an off peak plasmablast derived Ab, it may reflect a target of non-specific background plasmablasts that circulate at low percentages between period of antigen stimulation. Notably, these Abs that targeted KD4-2H4 had variable binding on prior published assays ^32^ so possibly our competition assay did not fully reflect optimal antigen targeting.

### Mimotope derived from Hepatitis C Virus (HCV)

Mimotope discovery is purely based on structural homology, so interpretation of specific peptides should proceed cautiously. We were using this study to specifically find a mimotope that may not have any biological releavance to the underlying condition. The KD4-2H4 targeting Abs may be similarly non-specific. Recent data suggests HCV is associated with autoimmune disorders ^33–35^ which may be related. Notably, 10076, the subject from which 76C group Abs was derived, was reportedly negative for HCV ^36^.

### Other Autoimmune targets of germline VH1-02 constructed Ab

Of the human protein targets found on the microarrays, autoAbs to these proteins have not been described in HIV. As many of these contain numerous acidic domains and relatively lower binding specificity in the initial peptide screen, these are likely non-specific reactions. AutoAbs to TMOD1 have been associated with pancreatic cancers ^37^ and IGA nephropathy ^38^ but no literature related to KD or HIV was discovered on our review. A number of the HIV bnAbs have been described as having autoimmune potential ^39,40^. Prior studies suggest gp41 targeting during initial infection relies predominantly on stimulating memory B cells that have previously been activated by non–HIV-1 antigens. A similar study reverting to germline other gp41 targeting Abs lost HIV reactivity but gained poly-reactive to various host or gut flora antigens ^41^ . Groups have postulated that germline Abs primed by reactions to commensal bacteria can be stimulated and form the basis for anti-gp41 Ab responses after infection ^42^. On our screen, the peptides showing highest binding were generally not derived from organisms that would fall into the ‘gut microbiome’ realm (see Table 1 and supplementary Table 1). It is possible that there is some microbiome dysregulation in both KD and HIV that may explain the cross-reactivity to KD4-2H4.

## Conclusion

Herein we identify a mimotope of a complex epitope that has been associated with functional Abs that associate with long-term non-progression. Since there have been no confirmatory studies supporting an association of HCV with KD, and we herein show no assocation of serum targeting in our KD samples, we believe this mimotope is a viable candidate to advance to pre-clinical HIV vaccination studies.

## Abbreviations

6F5: 76-Q13-6F5
6F11: 76-Q7-6F11
76Canc: 76C group ancestor
AA: amino acids
Ab/Abs: antibody/antibodies
ADCC: antibody dependent cell cytotoxicity
autoAbs: autoantibodies
bnAbs: broadly neutralizing antibodies
GLRX3: glutaredoxin 3
HCV: hepatitis C virus
HIV: Human Immunodeficiency Virus
IVIG: intravenous immunoglobulin
KD: Kawasaki Disease
MPER: membrane-proximal external region
TMOD1: Tropomodulin 1

## Acknowledgements

Study was supported by the Wildermuth Research Foundation (M.D.H) through the Variety Club of Buffalo and NIH R01 AI 125119-01 (M.D.H.); “The role of non-broadly neutralizing antibodies targeting gp41 structural epitopes in long term nonprogression of HIV infection.” M.D.H is a site PI for a Pfizer study, unrelated to the contents of this manuscript. Data available by request.

